# High fat diet initiates rapid adaptation of the intestine

**DOI:** 10.1101/2022.03.31.486473

**Authors:** Jacob R. Enriquez, Heather A. McCauley, Kevin X. Zhang, J. Guillermo Sanchez, Gregory T. Kalin, Richard A. Lang, James M. Wells

**Affiliations:** Division of Developmental Biology, Cincinnati Children’s Hospital Medical Center, Cincinnati, OH 45229-3039; Division of Endocrinology, Cincinnati Children’s Hospital Medical Center, Cincinnati, OH 45229-3039; Center for Stem Cell and Organoid Medicine (CuSTOM), Cincinnati Children’s Hospital Medical Center, Cincinnati, OH 45229-3039; Pluripotent Stem Cell Facility, Cincinnati Children’s Hospital Medical Center, Cincinnati, OH 45229-3039; The Visual Systems Group, Abrahamson Pediatric Eye Institute-Division of Pediatric Ophthalmology, Cincinnati Children’s Hospital Medical Center, Cincinnati, OH 45229-3039; Department of Ophthalmology, University of Cincinnati College of Medicine, Cincinnati, OH, USA; Medical Scientist Training Program, University of Cincinnati College of Medicine, Cincinnati, OH, USA

**Keywords:** fatty acid metabolism, intestinal stem cells, nutritional adaptation

## Abstract

While the systemic impacts of overnutrition are well-known to cause obesity and metabolic syndrome over the course of months, the immediate adaptive response of the intestinal epithelium to dietary changes remains poorly understood. Here we used physiological metrics and single cell analyses to interrogate the adaptive response of intestinal epithelial cells when moved to a high fat diet (HFD). Within 1 day of HFD exposure, mice exhibited altered feeding behavior, an increase in energy expenditure and an increase in intestinal epithelial proliferation. Single cell transcriptional analysis demonstrated several acute cellular changes on day 1, including a cell-stress response in intestinal crypts, de-granularization of Paneth cells, and a shift towards fatty acid cellular metabolism. By 3 days of HFD, there was an emergence of uncommitted progenitors with a transcriptional profile indicative of a shift from secretory populations and towards an absorptive fate. In enterocytes, genes regulating lipid transport and absorption increased over the first 3 days which paralleled a functional increase in lipid absorption *in vivo* over the course of 7 days on HFD. These findings demonstrate that intestinal epithelial cell populations respond rapidly to changes in diet through initial changes in cellular function followed by a shift in cellular composition.

## Introduction

From 1999 through 2018, the prevalence of obesity within the United States has increased from 30.5% to 42.4%^1^. In parallel, the prevalence of adult metabolic disease, including diabetes and dyslipidemia, increased from 25.3% to 34.2%, effectively making over one third of all US adults meeting criteria for metabolic syndrome^2^. These data highlight the urgency to understand metabolic disease for informed development of treatments. One factor that influences metabolic outcome is diet. The gastrointestinal tract (GIT) is the primary site for nutrient sensing and absorption of carbohydrates, proteins, and fats that are then utilized by cells of the body to maintain energy homeostasis. There have been numerous studies using animal models investigating the long-term impact of a high fat diet (HFD) on metabolic health^3,4,5,6^. For example, a single cell RNAseq analysis of mice on 12 weeks of a high fat/high sugar diet revealed intestinal maladaptation including altered intestinal lineage allocation^7^. However, by 12 weeks, animals are obese, have marked metabolic disease, and the intestine has undergone a significant adaptive response. Previous studies suggest that metabolic changes such as insulin resistance, hepatic steatosis, and ketogenesis may begin within days of HFD exposure^8,9,10,11,12^. However, it remains unknown how quickly the intestinal epithelium reacts and adapts to HFD, which is surprising given its essential role in systemic nutrient homeostasis.

The intestinal epithelium arises from proliferative intestinal stem cells (ISCs) that differentiate into specialized lineages along the crypt-villus axis^13^. ISCs first form secretory and absorptive progenitors that migrate through the transit-amplifying (TA) zone and gradually mature to become absorptive enterocytes or various secretory cells. Enterocytes comprise of the bulk of the intestine and absorb sugars, small peptides, and lipids. In parallel, secretory cells, such as Goblet, Paneth, or Enteroendocrine (EEC), secrete mucins, defensins, or hormone-peptides respectively in response to luminal stimuli. Recently, it has been shown that fasting, caloric restriction, and longterm HFD modulate ISC function and proliferation ^7,14,15,16^, suggesting that the intestinal epithelium is exquisitely sensitive to changes in diet. However, the immediate impacts of HFD on ISC differentiation into the various epithelial cells remains unknown.

Here, we sought to investigate the adaptive response of the different epithelial cell populations in the first week following a shift to HFD. We used whole body metabolic analyses and single-cell transcriptomics, combined with functional validation to identify the molecular basis of metabolic adaptation in the intestine. We observed immediate whole-body metabolic and intestinal crypt-niche stress after just 1 Day of HFD, an emergence of uncommitted progenitors fated towards enterocytes after 3 Days HFD, and functional adaptation to more efficient fatty acid uptake within 7 Days HFD. These data demonstrate that the intestine rapidly alters its transcriptional, cellular, and functional profile in response to changes in nutrient inputs.

## Results

### One day of HFD induces changes in mouse metabolism and IEC proliferation

While the long-term effects of a HFD on whole body metabolism has been studied for decades, the immediate impact of shifting to HFD is unknown. We used indirect calorimetry to monitor adult, wild-type mice maintained on Normal Chow (kcal% Fat-13%; Carbohydrate-58%; Protein-29%) or switched to HFD (kcal%: Fat-60%, Carbohydrate-20%, Protein-20%) for 7 Days (Figure 1A). We measured Oxygen Consumption (VO2) and Carbon-Dioxide Expiration (VCO_2_) (Figures 1B and 1C) to calculate the Respiratory Exchange Ratio (RER), which reflects the primary fuel source metabolized by the body. A ratio closer to 1.0 indicates predominantly glucose metabolism whereas a ratio closer to 0.7 suggests reliance on fatty acid metabolism^17^. Within the first day of HFD feeding, RER levels decreased from around 0.9 to 0.8 and this difference was maintained over time (Figure 1D). Moreover, we measured the Energy Expenditure (EE), which measures total energy required for homeostasis^18^. Within the first day, EE was increased from around 0.2-0.5 kcal/min to around 0.4-0.6 kcal/min (Figure 1E). We saw no significant changes in water intake or ambulatory movement over the 7 days (Figures S1A-S1C). We separately measured mouse weight, daily energy intake, and food intake from mice in normal housing. From the first day, mice given HFD gained weight and increased energy intake (Figures 1F and 1G). HFD food intake displayed significantly decreased slope over time (Figure 1H), suggesting that due to the higher caloric value of HFD, mice needed to consume less chow to achieve satiety. These data suggested that mice experienced whole-body metabolic changes within one day of eating a HFD, and that there may be related changes along the GIT.

**Figure 1.**
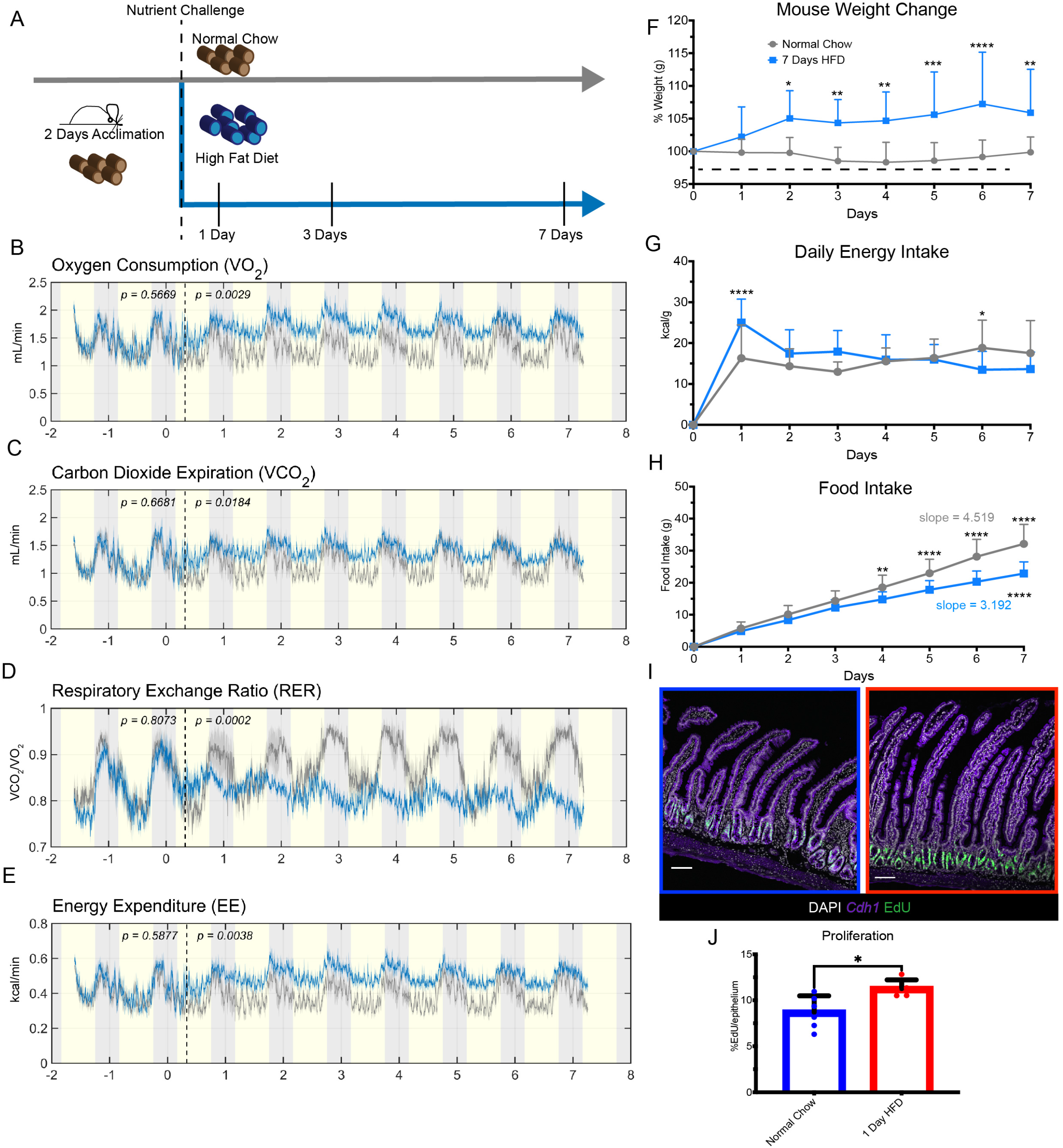
Mouse metabolism adapts to HFD within 1 day of exposure. (A) Diagram of feeding schedule in Metabolic Cages. Mice underwent 2 days of acclimation to the cages and all consumed Normal Chow (brown color pellets) during the acclimation period. The dashed line denotes when mice were divided into two groups, one maintained on Normal Chow (n=5), and the other group switched to High Fat Diet (HFD, n=6) for 7 days. (B) Average oxygen consumption (C) Average carbon dioxide expiration rate (D) Average respiratory exchange rate ratio calculated by VCO_2_/VO_2_ (E) Average energy expenditure measured in kcal/min captured from animals fed Normal Chow (Gray Line) and HFD (Blue Line). Yellow shading indicates light cycle whereas gray shading indicates night cycle. The dashed line indicates the end of the acclimation period and beginning of dietary challenge. (F) Percent daily weight change measured in grams (g), n= 11 Normal Chow, n= 20 HFD. (G) Average daily energy intake measured in kcal/g, n= 11 Normal Chow, n= 20 HFD. (H) Average accumulated food intake over time with calculated average slopes displayed, n= 11 Normal Chow, n= 20 HFD. (I) Immunofluorescence images of the proximal jejunum of mice fed Normal Chow (Blue Border) and 1 Day HFD (Red Border) after a 2-hour pulse with EdU to label actively proliferating cells (green). Nuclei are stained in white and *Cdh1* marks the epithelium in purple. Scale bars = 100μm. (J) Quantification of (I), n=6 Normal Chow, n=5 HFD. (B-E, J) Error bars are SD. One-Way ANOVA P-Values reported in panel. (F-H) Two-Way ANOVA; p-values: * < 0.05, ** < 0.0005, *** < 0.0003, **** < 0.0001

The proximal small intestine is the primary site of fat digestion and subsequent absorption; specifically, the duodenum is the site for lipase and bile acid activity to initiate emulsification, whereas the proximal jejunum absorbs the bulk of fatty acids^19^. Therefore, we investigated whether there was an adaptative response within the proximal intestine to acute HFD mediating these whole-body metabolic changes. Because the stem cell niche is sensitive to changes in dietary nutrients^14,15,16^, we examined intestinal proliferation, crypt depth, and villus height after 1 Day, 3 Days, or 7 Days of exposure to HFD. We only observed increased EdU incorporation after 1 Day of HFD and did not observe any changes in crypt depth or villus height (Figures 1I and 1J; Figures S1D-S1F). This suggested to us that HFD triggered an adaptive response within the intestinal epithelium within 1 Day, but that adaptative responses were likely to involve transcriptional or functional changes rather than growth in intestinal surface area.

### HFD initiates a rapid shift towards cellular fatty acid metabolism in intestinal lineages

To investigate how individual cell populations adapt to HFD, we performed single-cell RNA (sc-RNA) sequencing on sorted live, epithelial cells dissociated from the proximal intestine of adult mice (Figures S2A-S2C). Given its essential role in lipid absorption we largely focused on the proximal jejunum. After quality control and filtering of cells, we integrated the datasets of cells from mice eating Normal Chow, and those fed 1 Day HFD, 3 Days HFD, and 7 Days HFD and utilized the Normal Chow cells as the reference dataset^20^. This technique allowed us to not only determine the transcriptional heterogeneity present within each cluster, but also to compare transcriptional changes between conditions through a total number of 10,593 cells (Normal Chow: 2214, 1 Day HFD: 2309, 3 Days HFD: 2353, 7 Days HFD: 3717). Cell clustering analyses identified differentiated cell populations including Enterocytes, Enteroendocrine (EEC), Goblet, Tuft, and Paneth cells, as well as Enterocyte Progenitors (EP), Secretory Progenitors (SP), and Stem/Early-Transit Amplifying (TA) Zone cells (Figure 2A). The top 5 genes between each cluster verified known genes distinguishing between cell types (Figure 2B).

**Figure 2.**
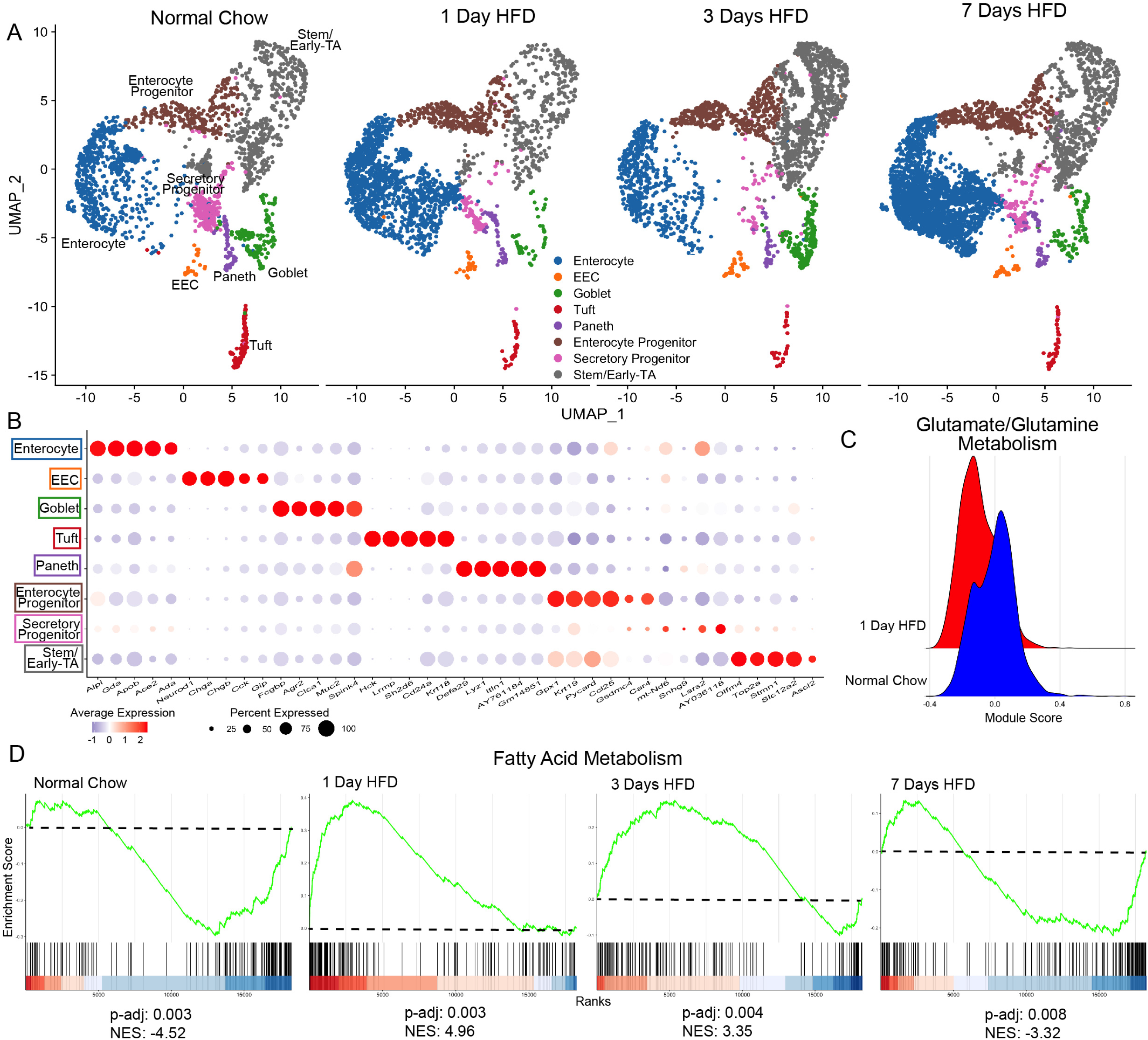
scRNA Seq Reveals Immediate shift towards Fatty Acid Metabolism. (A) Integrated analysis of jejunal epithelial sorted dataset-Normal Chow (reference dataset for integration, 2214 cells), 1 Day HFD (2309 cells), 3 Days HFD (2353 cells), 7 Days HFD (3717 cells). (B) Average expression of top 5 genes per cluster. (C) Ridge plot between Normal Chow and 1 Day HFD depicting module gene score calculated for glutamate/glutamine metabolism (D) Gene Set Enrichment Analysis (GSEA) using all clusters focusing on Hallmark Fatty Acid Metabolism genes compared between each dietary condition. Green line denotes enrichment score for each cell per condition. Dashed line marks score at 0.0. Black lines above color gradient denote localization of Hallmark-Fatty Acid Metabolism genes. Blue (low) to red (high) color gradient depicts enrichment score expression. Adjusted p-value (p-adj) and Normalized Enrichment Score (NES) reported in panel.

Analysis of the transcriptional changes in response to HFD using Biological Process Gene Ontology Terms (GO-Terms) suggested an increase in fatty acid metabolic pathways in several cell types at 1 Day HFD (Figures S2D-S2G) indicating a possible shift away from the normal glutamine/glutamate metabolism that is utilized by the intestinal epithelium^21^. To further investigate this we used genes collated by the Molecular Signatures Database (MSigDB) to evaluate the transcriptional signature of Glutamate/Glutamine Metabolism and saw an immediate downregulation after 1 Day of HFD (Figure 2C). Conversely, using gene set enrichment analysis (GSEA) we observed that most epithelial cells switch to utilizing fatty acid metabolism as indicated by normalized enrichment scores (NES) upon HFD exposure (Figure 2D). Specifically, NES for fatty acid metabolism showed an increase at both 1 Day and 3 Days HFD, suggesting an immediate metabolic adaptation to the influx of luminal fat. Interestingly, by 7 Days, NES for fatty acid metabolism decreased suggesting that enterocytes have adapted to the shift to HFD. This initial analysis revealed immediate metabolic transcriptional adaptations to the sudden change in nutritional input.

### HFD triggers an immediate stress response

Aside from immediate metabolic changes, we also observed an upregulation in genes associated with cellular stress (heat shock protein genes such as *Hspa1a*, *Hsp90aa1*, *and Hsf1*; unfolded protein response genes such as *Xbp1*, *Atf4*, *Eif2ak3*) for all epithelial cell populations after 1 Day HFD (Figure S3A). Stem/Early-TA and Paneth cells trended among the highest cellular stress responses (Figure 3A). Moreover, Stem/Early-TA cells revealed significant upregulation of genes encoding heat-shock related proteins, while Paneth cells displayed immediate downregulation of defensin related genes, suggesting an environmental stress response coupled with Paneth cell degranulation (Figure 3B and Figure 3C). To investigate this stress response and whether degranulation could be observed, we examined the ultrastructural level of intestinal tissue from mice via transmission electron microscopy (TEM). After 1 day on HFD we observed abnormal subcellular structure such as loss of granules and abnormal endoplasmic reticulum (ER) in Paneth cells (Figure 3D). Previous studies have shown that environmental or luminal factors such as smoking or Crohn’s Disease (CD) result in abnormal Paneth cell ultrastructure^22,23^. We also performed GSEA analysis using unfolded protein response (UPR) genes for Stem/Early-TA cells and inflammatory response genes for Paneth cells and saw significant increases in NES between Normal and 1 Day HFD for both subsets (Figures 3E and 3F). Heat shock proteins not only mediate the UPR by acting as protein chaperones, but also they initiate adaptation in response to inflammation^24^. These data suggest that cell stress within the stem cell niche is an early response to HFD prior to nutritional adaptation.

**Figure 3.**
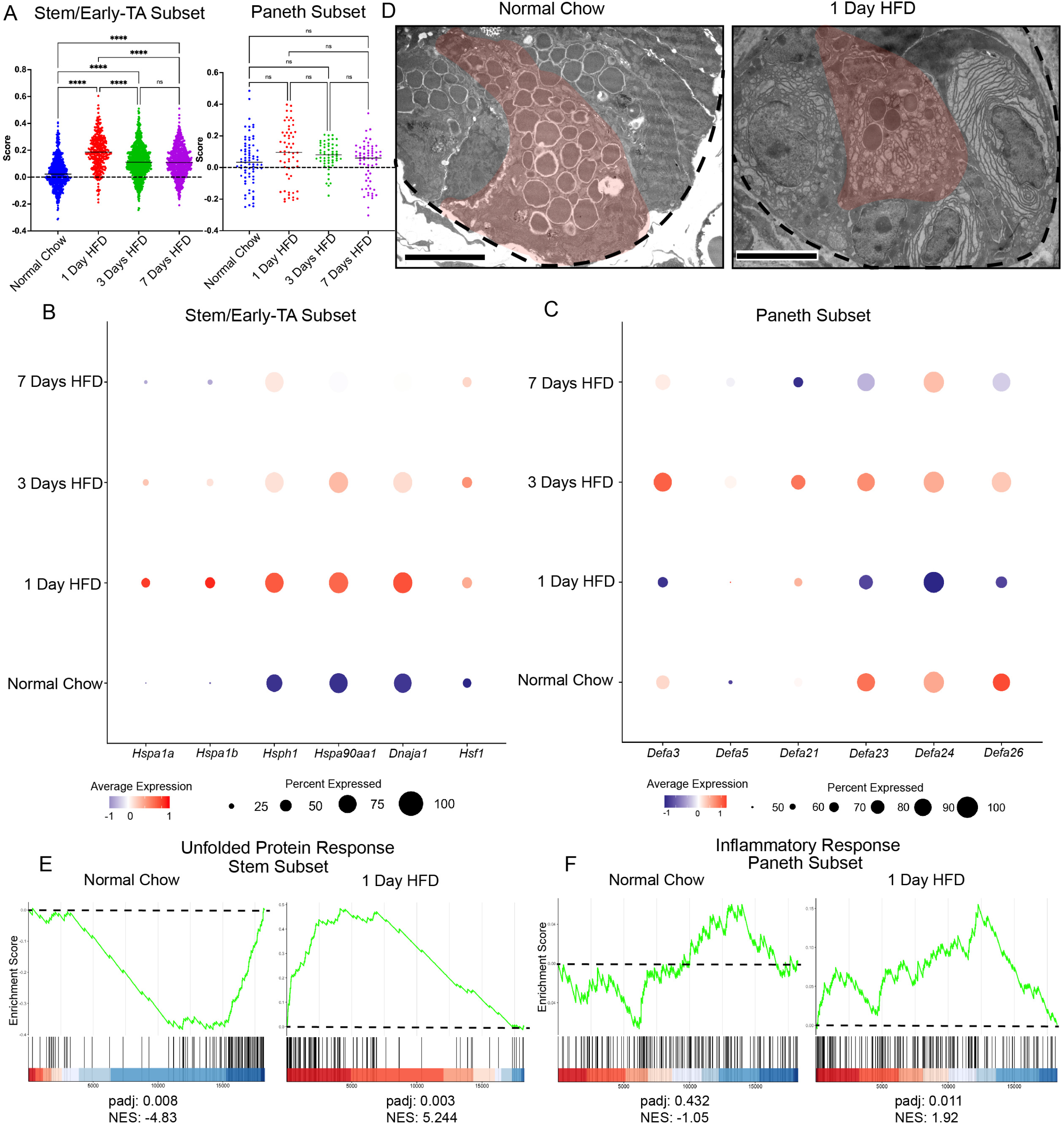
Intestinal crypts display cell stress immediately after exposure to HFD. (A) Cell stress module gene scores between dietary conditions in stem/early TA zone cells and in Paneth cells. (B) Dot plot depicting average gene expression of heat shock-related genes in Stem/Early TA-Zone cells. (C) Dot plot depicting average gene expression of genes involved in defensin regulation in Paneth cells. (D) Transmission Electron Microscopy (TEM) images of crypts from animals fed Normal Chow and 1 Day HFD. Dashed line outlines the bottom of the crypt while the red-pseudocolor denotes a representative Paneth cell. Scale bars = 4 μm. (E) Gene Set Enrichment Analysis (GSEA) of Hallmark Unfolded Protein Response genes comparing Stem/Early TA-Zone cells between Normal Chow and 1 Day HFD. (F) GSEA of Hallmark Inflammatory Response Genes comparing Paneth cells between Normal Chow and 1 Day HFD. Green line denotes enrichment score for each cell per condition. Dashed line marks score at 0.0. Black lines above color gradient denotes localization of Hallmark genes. Blue (low) to red (high) color gradient depicts enrichment score expression.

### Functional adaptation of enterocytes with improved lipid absorption and transport

Given that enterocytes mediate absorption of fat, we predicted that they would adapt to HFD. We calculated a lipid absorption score with known genes involved in fatty acid absorption. In comparison to the other cell types, enterocytes showed a pronounced increase in the lipid absorption score over time (Figure 4A; Figure S3B).

**Figure 4.**
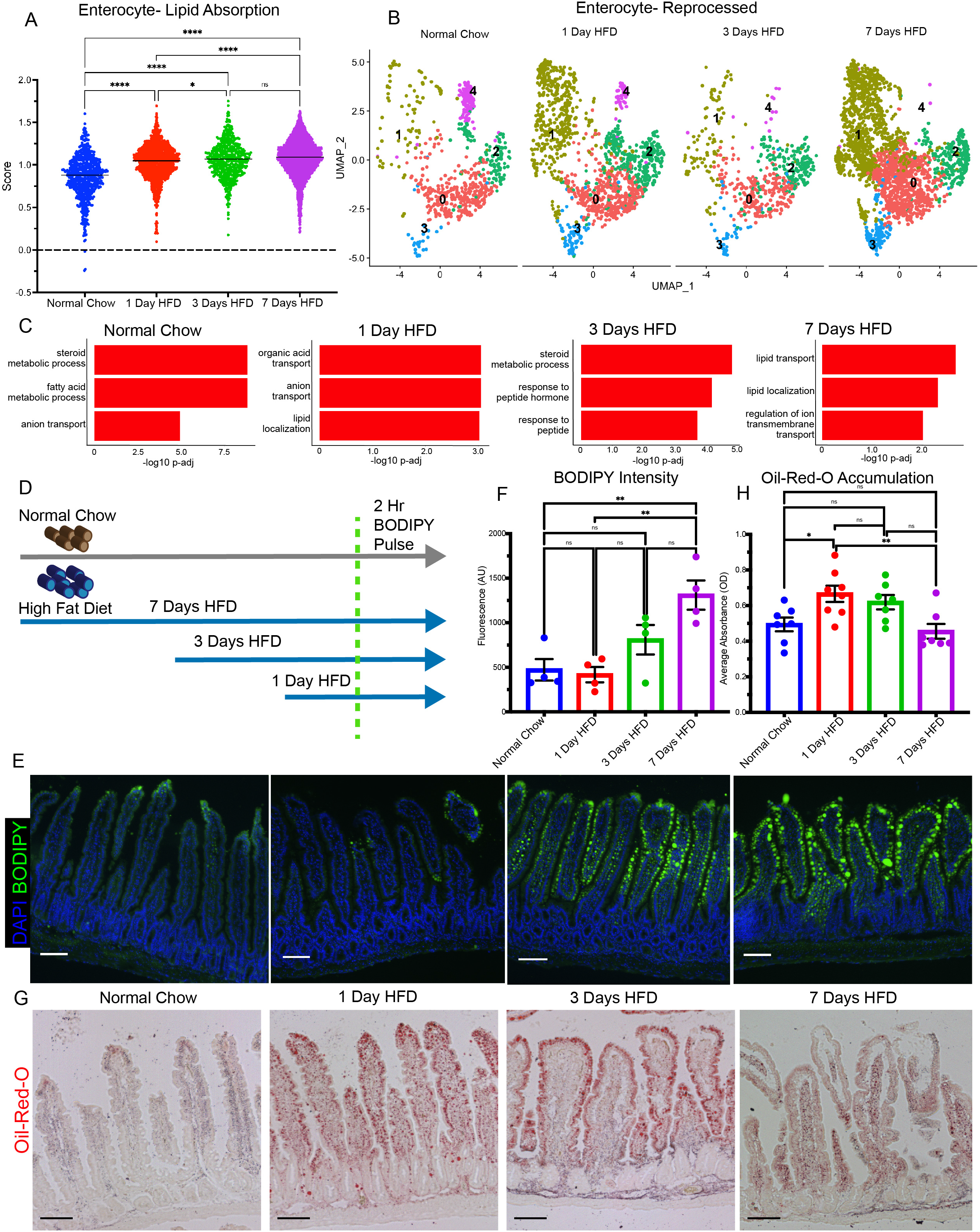
Enterocytes efficiently shift towards lipid absorption by 7 days HFD. (A) Lipid absorption score compared between enterocytes in each dietary condition. (B) Enterocytes were re-clustered into 5 subsets: 0 (1735 cells), 1 (1671 cells), 2 (847 cells), 3 (281 cells), 4 (200 cells). Breakdown per condition: Normal Chow (675 cells), 1 Day HFD (1308 cells), 3 Days HFD (549 cells), 7 Days HFD (2202 cells). (C) Top 3 Upregulated Gene Ontology Terms (Biological Process) for each condition in enterocyte subclusters 0 and 1. (D) Diagram of BODIPY Pulse-Chase Experiment. Normal Chow-gray arrow; HFD-blue arrow; green dashed line denotes time when all mice were gavaged with BODIPY and analyzed 2 hours later. (E) Fluorescence images of BODIPY (green) lipid absorbance in proximal jejunum 2 hours post-gavage. DAPI counterstains nuclei in blue. Scale bars = 100μm (F) Quantification of (E) fluorescence intensity. n=4 per condition. (G) Oil-Red-O staining of proximal jejunum for each condition. Scale bars = 100μm (H) Quantification of (G) reported as optical density (OD) for Oil-Red-O accumulation. n=7 for each condition. (A, F, H) Error bars are SEM. One-Way ANOVA; p-values: ns-not significant, * < 0.05, ** < 0.005, **** < 0.0001

Interestingly, the other cell clusters (EEC, Paneth, Tuft, EP, SP, Goblet, and Stem/Early-TA) also experienced a small increase in score at 1 Day HFD, possibly due to the shift to fatty acid metabolism as a primary energy source. To determine whether specific enterocytes were responding to HFD, we specifically re-analyzed the enterocyte cluster and compared GO-Term Biological Process analysis between each new subcluster (Figure 4B). This analysis revealed 5 new enterocyte sub-clusters; where sub-cluster-0 and −1 displayed upregulated transport related processes (Figure S3C). These data support previously published specialized functions of enterocytes along the crypt-villus axis^16^. Furthermore, by specifically focusing on enterocytes regulating transport, we could transcriptionally analyze the response of these enterocyte sub-clusters to HFD. Therefore, with sub-cluster-0 and −1, we performed GO-Term Biological Process analysis and the top 3 terms between each condition revealed a shift towards lipid transport, lipid localization, and regulation of ion transmembrane transport by 7 Days HFD (Figure 4C). Intestinal uptake of fatty acids is mediated via passive diffusion or through active transporters, such as *Cd36* or *Fatp4* (*Slc27a4*)^25,26^. Enterocytes in subclusters 0 and 1, showed a significant increase in *Slc27a4*, with some increase in *Cd36*(Figures S3D-S3E), indicating a shift from passive to active transport of fatty acids. Together these data suggested the emergence of a newly adapted population of enterocytes that are transcriptionally specialized towards efficient transport of lipids.

To assess whether these transcriptional changes resulted in increased efficiency of lipid absorption, we analyzed enterocytes in the proximal jejunum in two ways. First, we took animals that had been on HFD for 1, 3, and 7 days and gavaged them with a bolus of fluorescent-labeled fatty acid BODIPY dye (Figure 4D). We then isolated the jejunum after 2 hours and quantified the uptake of lipids by enterocytes. When compared to Normal Chow, enterocytes adapting to HFD displayed markedly increased capability of absorbing fatty acids, with efficiency increasing most at days 3 and 7 (Figures 4E and 4F). Using a separate approach, we visualized lipid droplet accumulation using Oil-Red-O, which measures the steady-state presence of lipids in cells (Figure 4G). In enterocytes from animals maintained on Normal Chow, lipid accumulation was observed at the villus tips with some diffuse staining in the lamina propria. In response to a shift to HFD, staining at day 1 was the highest with lipids accumulating along the whole villus length and beginning to accumulate in the lamina propria (Figure 4H). At day 3 of HFD, lipid accumulation in enterocytes began to diminish, but increased in the lamina propria. At day 7, the jejunum seems to have adapted to HFD, with lipid accumulation being similar to Normal Chow. This change in droplet localization coupled with efficient fatty acid uptake led us to conclude that there was an increase in lipid transport capability by enterocytes mediated by transcriptional adaptation. These data demonstrate that enterocytes transcriptionally and functionally shifted towards efficient lipid absorption within days of a dietary switch to HFD.

### Lineage tracing, sorting, and single cell RNAseq analysis of rare epithelial cell types

Absorptive enterocytes are the most abundant cell population in the intestine. Some secretory lineages, particularly EEC and Paneth cells are relatively rare. Within our proximal jejunal dataset (Figure 1), we only captured 158 total cells (1.5% of total cells). However, we were interested to determine the impact of acute HFD on EEC populations because EECs are known to secrete hormones in response to luminal stimuli, including fatty acids^27^. To increase our power analysis of rare secretory cells, we performed genetic cell labeling and sorting using *Neurog3* Cre-Recombinase and a tdTomato reporter. Previous studies demonstrated that *Neurog3-Cre* labels EECs, as well as multiple non-endocrine cell lineages including progenitors, and we employed this as a means to enrich for these rare populations ^28,29^. We sorted live, epithelial, tdTomato+ cells from *Neurog3Cre-tdTom* mice consuming Normal Chow, 1 Day HFD, 3 Days HFD, or 7 Days HFD and performed sc-RNA sequencing on the proximal intestine with the same parameters as our initial dataset (Figures S4A-S4B). As expected, we enriched for EECs (4915 total cells, 23% of total cells), as well as other non-endocrine cell types such as Paneth, Goblet, and even Stem cells (Figures S4C-S4E). We confirmed this analysis via immunofluorescence staining where we saw rare lineage traced ribbons emanating from the crypts, while also individual cell type marker genes co-localized with tdTomato in cells independent of crypts (Figure S4F-S4G). A comparison of unenriched Epithelial only cells, TdTom+ enriched, and a previously published single cell dataset^30^ determined that TdTom+ sorted populations not only contained all expected cell populations, but also substantially enriched for secretory progenitor and secretory populations (Figures S4H-S4K).

With this highly powered data set, we first analyzed the transcriptional response of the EEC population to short-term HFD. There were 9 different clusters segregating into known EEC-hormone subtypes (Figures S5A-S5B) identified by expression of Glucagon (*Gcg*), Peptide-yy (*Pyy*), Gastric Inhibitory Polypeptide (*Gip*), and Cholecystokinin (*Cck*), which are known hormones released in response to luminal fatty ingestion ^31,32^. Given their role in nutrient sensing, we predicted that EEC transcription would be highly impacted by HFD. Surprisingly, transcription of EEC hormones in the proximal intestine modestly changed, for example transcription of *Gip* increased over 7 Days HFD, whereas *Gcg*, *Pyy*, and *Cck* expression decreased slightly at 3 Days HFD before increasing back up by 7 Days HFD (Figure S5C). We confirmed this analysis by performing GIP and CCK ELISAs on mouse serum for all time points and saw an increasing trend of GIP by 7 Days HFD, whereas CCK levels slightly decreased at 3 Days HFD but remained relatively constant (Figure S5D). Moreover, expression of known fatty acid receptors, such as Ffar1, Ffar2, Ffar3, Ffar4, and Gpr119^33^ remained relatively unchanged over the course of HFD (Figure S5E). Overall, EEC hormones and EEC fatty acid receptor transcriptional activity remained resilient during short term HFD and were not easily manipulated by fatty acid exposure.

### HFD causes a shift in cell lineage allocation from secretory to absorptive cell types

In addition to the immediate transcriptional response of ISCs to high fat diet at day 1, we also observed a reduction of all secretory lineages and an increase in enterocytes over the 7 days of HFD in the proximal intestine. In unenriched epithelial cell populations, Paneth, EEC, Goblet, and Secretory Progenitor populations were all reduced in numbers during 3 days of HFD (Figure S2B) whereas enterocyte progenitors and enterocytes were all increased over the same time frame (Figure 5C). This suggested that, in response to HFD, ISCs shifted their differentiation preference, from secretory to absorptive. Since secretory lineages, like EEC and Paneth cells are rare, this initial conclusion was based on a relatively small number of cells. We therefore analyzed the secretory cell-enriched dataset to determine any lineage allocation shifts.

**Figure 5.**
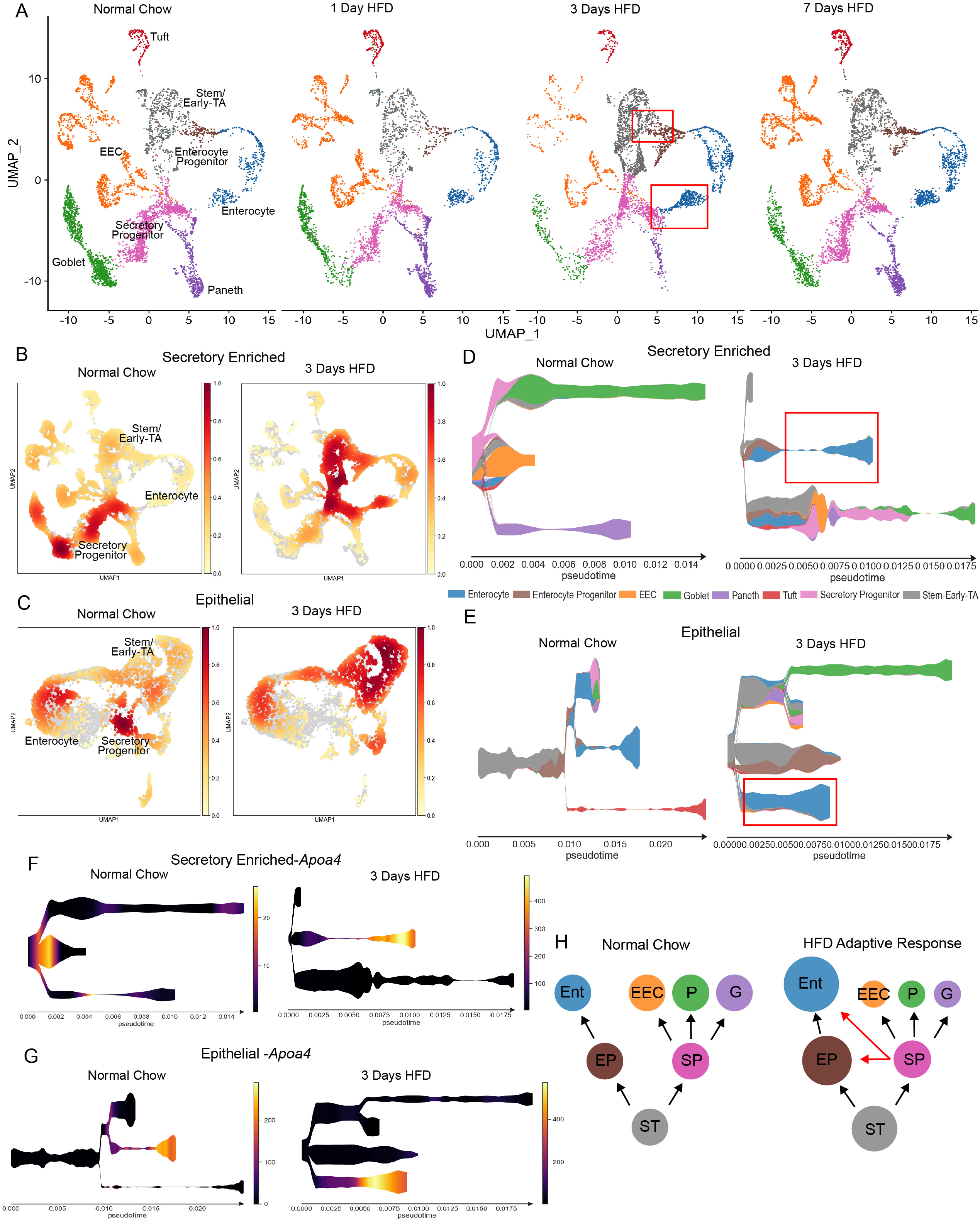
Emergence of progenitors computationally fated for enterocytes at 3 days HFD. (A) Integrated reference analysis for secretory enriched (sorted TdTom+) cells for each condition colored by each cluster. Red box at 3 Days HFD focuses on emerging cluster of new cells categorized within Stem/Early-TA and Secretory Progenitor bridging between Enterocyte Progenitor and Enterocyte clusters. (B) Distribution of Secretory Enriched cells depicted as density plot split between mice eating Normal Chow and 3 Days HFD. (C) Distribution of epithelial cells (non-labeled dataset) depicted as density plot split between mice eating Normal Chow and 3 Days HFD. Yellow (low) to red (high) color gradient depicts density value. (D) STREAM pseudotime analysis for Secretory Enriched cells from mice eating Normal Chow and 3 Days HFD. Stem cluster was used as the origination node and then unbiased analysis of clustering with 3 branches. (E) STREAM pseudotime analysis for epithelial cells from mice eating Normal Chow and 3 Days HFD. Stem cluster was used as the origination node and then unbiased analysis of clustering with 5 branches. (F) STREAM River plots of *Apoa4* (Apoliprotein-A4) distinguishing between Normal Chow and 3 Days HFD for Secretory Enriched cells. Y-axes are log-scale expression values. (G) STREAM River plots of *Apoa4* distinguishing between Normal Chow and 3 Days HFD for Epithelial dataset. Y-axes are log-scale expression values. (H) Diagram delineating intestinal epithelial cell differentiation priority during Normal Chow and the adaptive response to HFD (ST-Stem/Early TA-Zone, EP-Enterocyte Progenitor, SP-Secretory Progenitor, Ent-Enterocyte, EEC-Enteroendocrine Cell, P-Paneth, G-Goblet). Circle size denotes relative cell abundance. Red arrows during HFD indicate shift of cells towards Enterocyte Progenitors and Enterocytes.

(Figures S4C). We again observed an overall reduction in secretory cells, however the biggest reduction seemed to be after 3 days of exposure to HFD, where EECs, goblet and Paneth cells were all substantially reduced and there was a change in cell cluster density allocation (Figure 5A and Figure 5B). Also, at 3 Days HFD there was an increase in progenitor populations, including a new cluster of cells branching from the Stem/Early-TA and Secretory Progenitor clusters towards the Enterocyte clusters (Figure 5A).

We considered the possibility that these progenitor cells could represent the diversion of stem and secretory progenitors towards the enterocyte lineage. We therefore performed pseudotime analysis of all cells at 3 Days HFD and compared them to Normal Chow cells by using STREAM^34^. By setting the Stem Cell/Early-TA cluster as the origination node, the analysis revealed distinguishing trajectories between Normal Chow and 3 Days HFD (Figure 5D). As expected, the Normal Chow secretory enriched dataset delineated mainly secretory populations such as EEC, Goblet, and Paneth. However, at the 3 Days HFD timepoint, the Enterocyte Progenitor and Enterocyte lineages became uniquely distinct suggesting the shift in differentiation towards an absorptive lineage in response to HFD. Furthermore, this notion of lineage allocation trajectory was supported by performing similar analysis within the overall epithelial single-cell dataset (Figure 5E). We visualized this using one example of an Enterocyte-specific gene, apolipoprotein-A4 (*Apoa4*), which depicted a difference in enterocyte lineage allocation in Normal Chow versus 3 Days HFD (Figures 5F and 5G). Therefore, using two different single cell datasets and computational modeling, our data suggests that within one week of HFD exposure, the intestine restructures itself to better absorb dietary fats by preferentially diverting progenitor populations towards the absorptive lineage (Figure 5H).

## Discussion

This study revealed that immediately upon dietary switch from a normal/balanced diet to HFD, multiple cell types of the intestine coordinate an adaptive, stepwise response towards efficient fatty acid uptake. This is the first report investigating early mechanisms by which the intestinal epithelium responds and adapts to a change in diet. Within 24 hours, cellular populations within the crypt exhibited cell stress at the transcriptional and ultrastructural level. Over the next few days, there was an increase in cellular populations which we computationally projected to be biased toward enterocyte progenitor fate. This culminated in an altered intestinal epithelium adapted to highly efficient absorption of lipids by one week of HFD.

Previous studies using HFD to elicit metabolic phenotypes in animals overlook this initial window of adaptation. Most protocols expose mice to HFD for 6 weeks to 6 months^3,4,5,6^. Within the intestine over this prolonged HFD, the epithelial lining thins, secretory lineage abundances alter, and consequential morbidities of obesity and diabetes develop. Our data suggest that the intestine may acutely respond each time there is a major shift in nutrient input and that changes are immediately observed. Moreover, while the diets of laboratory animals are easily manipulated, the nutritional intake of humans is difficult to regulate. However, the ability of mammals to adapt to changing availability of nutrients is essential, especially with seasonal changes in diet.

Further studies are required to elucidate the benefits, and indeed harm, of short-term dietary changes humans experience.

Our study also suggests that cellular stress responses in the crypt to HFD precedes a change in differentiation trajectory of intestinal stem and progenitor cells. Previously, it has been shown that premature induction of endoplasmic reticulum (ER) stress and the unfolded protein response (UPR) causes loss of stem cell activity and expansion of transit-amplifying cells^35^. Moreover, ER stress and UPR are highly associated with intestinal diseases including inflammatory bowel disease (IBD) and colorectal cancer (CRC)^36^. Therefore, the repeated stress of being exposed to periods of HFD, followed by return to a more balanced diet, could mirror the effects of periodic inflammation. It may be possible that intermittent and/or prolonged cell stress upon every major dietary shift could induce these widespread gastrointestinal diseases.

This study also highlights the inherent plasticity of the intestinal epithelium. Here, we found that HFD stimulated intestinal proliferation within 24 hours, we observed a shift of progenitors from secretory cells to absorptive enterocytes, and that those enterocytes were functionally better equipped to absorb lipids by 7 days after the switch to HFD. Our data suggest that by one week of HFD, the intestine had adapted to maximize nutrient metabolism. Previous studies have described changes in intestinal stem cell function in response to a dietary change ^7,14,15,16^, however this study is the first to comprehensively describe the coordinated and stepwise progression of acute adaptation of multiple epithelial cell populations. Moreover, our data support the notion of stem and progenitor cell expansion in response to diet, similar to the many reports describing dedifferentiation into a stem/progenitor cell-like state in response to environmental stressors or injury ^37,38,39,40,41,42^. The pseudotime analysis showing the emergence of a new population of progenitors at 3 days of HFD suggest that altered lineage allocation is an adaptive response to dietary changes. Recently, another group performed single-cell transcriptomic analysis of mice fed a high sugar/high fat diet for 12 weeks and also observed dietary induced changes in intestinal stem cells and progenitor hyperproliferation^7^. However, it was not appreciated that adaptation happens within the first week. Taken together, our findings demonstrate the intestine undergoes a rapid and dynamic response to adjust to the changing availability of nutrients.

In summary, we took a multi-pronged approach to identify the initial steps of the adaptive response to HFD, including whole body metabolism, tissue functional and morphologic changes, as well as single cell transcriptomics. These transcriptional and functional analysis revealed immediate changes in all intestinal epithelial cell types occurring within 24 hours and resulting in a modified intestinal epithelium adapted to maximize absorption of fat within one week. This type of intestinal plasticity may be an evolutionary adaptation to periods of nutrient scarcity, but in current times of nutrient excess, could be contributing to rising rates of obesity, metabolic disease, and inflammation.

## Acknowledgments

We thank Dr. Kelli VanDussen, Dan Schnell, Jay Stone, and Praneet Chaturvedi for valuable guidance and discussion. We also thank support provided by the Confocal Imaging Core, Research Flow Cytometry Core, and Gene Expression Core at CCHMC. We thank the members of the Wells and Zorn lab for reagents and feedback. This research was supported by the grants from the NIH, U19 AI116491 (JMW), P01 HD093363 (JMW), UH3 DK119982 (JMW), the Shipley Foundation (JMW), and the Allen Foundation (JMW). We also received support from the Digestive Disease Research Center (P30 DK078392).

## Author Contributions

JMW and JRE conceived the study and constructed manuscript. JRE collected all biological samples, performed scRNA-seq experiments, computational analysis, and functional analysis. HAM and JGS aided in epithelial dissociation. GTK aided in EdU and immunofluorescence analysis. KXZ and RAL performed metabolic cage experiments and analysis. All authors approved of manuscript

## Declaration of Interests

The authors declare no competing interests.

## Methods

### Resource Availability

#### Lead Contact

Further information and requests may be directed to the lead contact, James M. Wells

#### Materials Availability

This study did not generate any new unique reagents.

#### Data and Code Availability

The data in this publication are currently being deposited in the Gene Expression Omnibus

### Experimental Model and Subject Details

All animal procedures were approved by the Cincinnati Children’s Hospital Research Foundation Institutional Animal Care and Use Committee (IACUC2019-0006) and performed using standard procedures. C57BL6/J and *Neurog3Cre* x Rosa26tdTom mice under BL6/C57 background were maintained on a 14L: 10D light cycle and had ad libitum access to chow and water. Normal Chow diet (NCD) administered (5010, LabDiet). For the acute high fat diet conditions, mice were given similar food weight of high fat diet (HFD, D12492, Research Diets, Inc.) in the early mornings 1 day, 3 days, and 7 days before time of experimental collection. Mouse weight and diet weight were weighed every morning. All mice were aged 9-14 weeks and both males and females were used in the study.

### Method Details

#### Indirect Calorimetry

9-14-week-old male C57BL6/J mice were acclimated in metabolic chambers (Promethion, Sable Systems International) for 2 days prior to the start of the study. Mice were continuously recorded for a total of 9 days at ambient room temperature (22°C) with the following measurements taken every 5 minutes: gas exchange (VO_2_ and VCO_2_), food intake, water intake, and spontaneous locomotor activity (cm s^-1^) in the XY plane. All animals were initially fed standard chow until day 2 of the study, when one group was switched to a 60 kcal% high-fat diet (HFD, D12492, Research Diets, Inc.), and one group remained on a standard normal-chow diet (NCD). Otherwise, food and water were available *ad libitum*. All mice were subjected to a 14L:10D light cycle for the duration of the acclimatization and study period. VO_2_, VCO_2_, and energy expenditure (EE) were calculated according to the manufacturer’s guidelines (Metascreen software v.2.3.15.11, Sable Systems International), with energy expenditure estimated via the abbreviated Weir formula. Respiratory exchange ratio (RER) was calculated by the ratio VCO_2_/VO_2_. Mass dependent variables (VO_2_, VCO_2_, energy expenditure) were not normalized to body weight. Food and water intake were measured by top-fixed load cell sensors, from which food and water containers were suspended into the sealed cage environment. For food consumption, mice demonstrating excessive food grinding behavior were excluded from statistical analyses. Raw data were exported using Sable Systems International ExpeData software v.1.9.27. Published data represent a combination of two independent experiments conducted separately. *P* values are from one-way repeated measures ANOVA comparing HFD and NCD groups before and after the introduction of the high-fat diet on study day 2.

#### Epithelial Single Cell Dissociation and FACS Sorting

Approximately 5-7 cms of duodenal intestine was dissected and then subsequently after the duodenal-jejunal flexure another 5-7 cms of proximal jejunal intestine was dissected for all dietary conditions. Each piece was filleted open to expose crypts and villi and rinsed with PBS before digested into single cells with 5 mLs Tryple and 5 uls Roc Inhibitor. Single cells were filtered through a 40 um cell strainer and then recorded using LSR Fortessa flow cytometer and analyzed with FACSDiva software. In all experiments, samples were labeled with Anti-EpCam-APC (BD Biosciences) to delineate epithelial cells and stained with SYTOX Blue dead cell stain (Life Technologies). The forward and side scatter plots were used to discriminate doubles and cellular debris. The secretory enriched datasets were additionally sorted for TdTom+ fluorescence.

#### Single-Cell RNA Sequencing Pipeline and Analysis

Single cell RNA-Sequencing (scRNA-seq) library preparation was performed by the CCHMC Gene Expression Core using the Chromium 3’v3 GEM Kit (10x Genomics, CG000183RevC). Approximately 12,800 cells were loaded to achieve 8,000 captured cells per sample to be sequenced. Sequencing was performed by the CCHMC DNA Sequencing core using the NovaSeq 6000 (Illumina) sequencing platform with an S4 flow cell to obtain approximately 320 million reads per sample. Raw scRNA-seq data was converted to FASTQ files and then aligned to the mouse genome [mm10] using default parameters of CellRanger (v3.0.2, 10x Genomics). Reads were aligned to mouse genome [mm10]. Quality control and clustering were performed using Seurat^20^ [v3.2.3] in R. Basic filtering parameters included cells with unique features of minimum 100 and maximum 7500. Cells expressing less than 50 percent mitochondrial related genes were included. Cell cycle effect was regressed out using previously established methods in Seurat. After initial filtering, each individual dataset was run through DoubletDecon^43^ using default parameters. Rhop values for each sample are as follows: Epithelial Datasets-Normal Chow: Duo-1 & Jej-1.2; 1 Day HFD: Duo-1.1 & Jej-1.2; 3 Days HFD: Duo-1.2 & Jej 1.1; 7 Days HFD: Duo-1.2 & Jej 1.1; Secretory Enriched Datasets-Normal Chow: Duo-1.2 & Jej-1.2; 1 Day HFD: Duo-1.2 & Jej-1.2; 3 Days HFD: Duo-1.2 & Jej-1.1; 7 Days HFD: Duo-1.2 & Jej-1.1. After identification of doublets and removal from each dataset, the newly filtered datasets were subsequently used for all downstream analysis. The Epithelial dataset samples were integrated using standard SCTransform Seurat protocols with Normal Chow Condition as Reference Dataset using 2000 integration features. Likewise, the Secretory Enriched Datasets were also integrated using standard SCTransform Seurat protocols with Normal Chow Condition as Reference Dataset using 2000 integration features. After clustering, cells were visualized using UMAP^44^. Marker genes were determined using ‘FindAllMarkers’ function (Wilcoxon rank-sum test) in Seurat. The initial Integrated Epithelial dataset showed some contaminating mesenchymal cells that were not removed during the sort. These cells were distinct from the other clusters and were subsequently removed from the final analysis.

To perform analysis in comparison to Haber *et al*., 2017 related to Figure S4, first we downloaded the Regional_UMIcounts.txt file from GSE92332. We then performed the standard SCTransform Seurat pipeline and subsequently filtered out only the Jejunum. We then performed SCTransform Seurat Integration between Normal Chow jejunal Epithelial, Normal Chow jejunal Secretory Enriched, and the Haber dataset with the Haber dataset as the Reference Dataset using 2000 integration features. Subsequent clustering and marker gene expression was performed as previously described.

To perform GSEA analysis, we used the msigdbr package (which pulls from MSigDB Collections) to determine lists of genes from the Hallmark Gene Sets (H: Fatty Acid Metabolism, Unfolded Protein Response, and Inflammatory Response). We then used presto to determine differentially expressed genes between conditions and then performed gene set enrichment anaylsis using fgsea.

To determine Glutamate/Glutamine Metabolism score, the integrated Epithelial dataset was used with the default setting for AddModuleScore() function of Seurat. We pulled genes from MSigDB Curated Gene Sets (C2: Reactome-Glutamate/Glutamine Metabolism). We then plotted results using RidgePlot() function of Seurat.

To determine Cell Stress Signature Score, the integrated Epithelial dataset was used with the default settings for AddModuleScore() function of Seurat. The following genes were used: Mgat4a, Dgat1, Dgat2, Npc1l1, Abcg5, Abcg8, Apoa1, Apoa4, Mttp, Slc27a4, Apob, Apoc3, Fabp6, Fabp2, Plin3, Sar1b, Cd36.

To determine Lipid Absorption Signature Score, the integrated Epithelial dataset was used with the default settings for AddModuleScore() function of Seurat. The following genes were used: Mgat4a, Dgat1, Dgat2, Npc1l1, Abcg5, Abcg8, Apoa1, Apoa4, Mttp, Slc27a4, Apob, Apoc3, Fabp6, Fabp2, Plin3, Sar1b, Cd36.

To visualize cell density plots, both the integrated Epithelial dataset and the Secretory Enriched dataset was migrated from Seurat to Scanpy^45^. Once data was formatted correctly in Scanpy, default settings for sc.tl.embedding_density were used to calculate Gaussian kernel density for all cells. Plots were generated by using sc.pl.embedding_density grouped by condition.

Pseudotime analysis was performed with default parameters of STREAM^46^. Visualization of river plots was also performed using st.plot_stream as log normalized gene expression.

#### Tissue Harvest and Immunofluorescence Imaging

Mouse proximal intestines were harvested and prepared for ‘swiss-roll technique’ to be fixed in 4% paraformaldehyde (PFA) at 4C overnight. The fixed samples were washed with PBS for 10 mins three times. The intestines were then immersed in 30% sucrose/PBS for 48 hrs. Afterwards, tissue sections were prepped by embedding each region in OCT and cryosectioned (8um) onto Superfrost Plus slides.

Immunofluorescence staining was performed by treating PBS re-hydrated tissue slides with 0.1% Triton X-100/PBS and then incubated with 5% normal donkey serum. Tissues were incubated overnight at 4C with the following primary antibodies: ECAD (rat; R&D 1:1000), Ace-2 (gt, 1:500), Lysozyme (Dako, rab, 1:200), Chga (ImmunoStar, rab, 1:200), Olfm4 (CellSignaling, rb, 1:200), Ckrt18 (Abcam, biotin, 1:200), Dclk1 (Novus Biologicals, rab,1:200). The next day, slides were washed with PBS for 10 mins three times, and then incubated with fluorescently conjugated secondary antibodies for the prospective animal. Nikon Confocal Fluorescent Microscopes were used for image capture.

#### Crypt Depth/Villus Height Measurements

Processed and imaged mouse proximal intestine was measured using NIS Elements by two independent viewers with blinded samples. Per mouse, between 70 and 150 crypts and villi were counted. Crypt depth was measured from the base of the crypt to the top of the transit-amplifying zone. Villus height was measured from the top of the transit-amplifying zone to the tip of a villus. Average values for each mouse were then plotted for each condition.

#### EdU-Cell Proliferation

On the day of collection after nutrient challenge, 9-14-week-old male and female mice were intraperitoneally injected with 100 mg/kg of 5-Ethynyl-2’-deoxyuridine (EdU). After waiting for 2 hours, intestines were collected for ‘swiss-roll technique’ as described in previous section. The standard protocol for the Click-iT™ EdU Cell Proliferation Kit (647) was then followed and subsequently imaged.

#### Oil-Red-O Staining

Oil-Red-O Stock (ScienCell, 0843a) was diluted 3:2 using deionized H2O to make a working solution. The working solution was then filtered with 0.22 um syringe driven filter unit (Millipore). Specimens, similarly prepared for immunofluorescence imaging on glass slides, were rehydrated in H2O for 5 mins, subsequently washed in 60% ethanol, and then exposed to filtered working Oil-Red-O solution for 20 minutes. After another 60% ethanol rinse, slides were rehydrated with H2O and then cover-slipped with Fluoromount-G. Imaging was performed on Nikon Widefield Fluorescent Scopes. To perform quantification, slides were processed the same but were not rehydrated; rather eluted with 100% ethanol for 1 hour. Then eluted dye from slides were transferred onto a 96-well imaging plate and absorption was measured in triplicate at 510 nm using a plate reader.

#### Transmission Electron Microscopy (TEM)

Proximal jejunum was dissected and fixed in 3% glutaraldehyde and 0.175 M sodium cacodylate buffer, pH 7.4, at 4°C for one hour. The samples were then post fixed in 1% osmium tetroxide in 0.2 M sodium cacodylate buffer for 1 hour at 4°C, processed through a graded series of alcohols, infiltrated, and embedded in LX-112 resin. After polymerization at 60°C for three days, ultrathin sections (100 nm) were cut using a Leica EM UC7 microtome and counterstained in 2% aqueous uranyl acetate and Reynolds lead citrate. Images were taken with a transmission electron microscope (TEM, Hitachi H-6750) equipped with a digital camera (AMT 2k×2K tem CCD).

#### BODIPY Gavage

Mice 9-14 weeks were placed on dietary challenge (Normal Chow or 1, 3, 7 Days HFD). On the final day of diet, mice were then gavaged with BODIPY (50 ug/mg) and swiss-rolled intestines were flash-frozen harvested 2 hours later. Sections were fixed with 4% PFA for 5 minutes and then stained with DAPI for 10 min. Slides were imaged using Nikon Upright Microscope.

#### Quantification and Statistical Analysis

For all confocal quantification, such as EdU and BODIPY analysis, each dot represents an individual “N” number of mice shown as reported in legend. For Indirect Calorimetry measurements, *P* values are from one-way repeated measures ANOVA comparing HFD and NCD groups before and after the introduction of the high-fat diet on study day 2. For mouse weight changes, daily energy intake, and food intake, *P* values are from two-way ANOVA Sidak’s multiple comparisons test. For comparison between conditions of module gene scores, *P* values are from one-way repeated measures ANOVA Tukey’s multiple comparisons test. Statistical analysis was performed in Prism 9 [v9.3.1].

## Notes

### Competing Interest Statement

The authors have declared no competing interest.

