## Supplementary Figures for "High fat diet initiates rapid adaptation of the intestine"

Figure S1- Additional metabolic cage information and morphometric analysis of proximal intestine during acute HFD

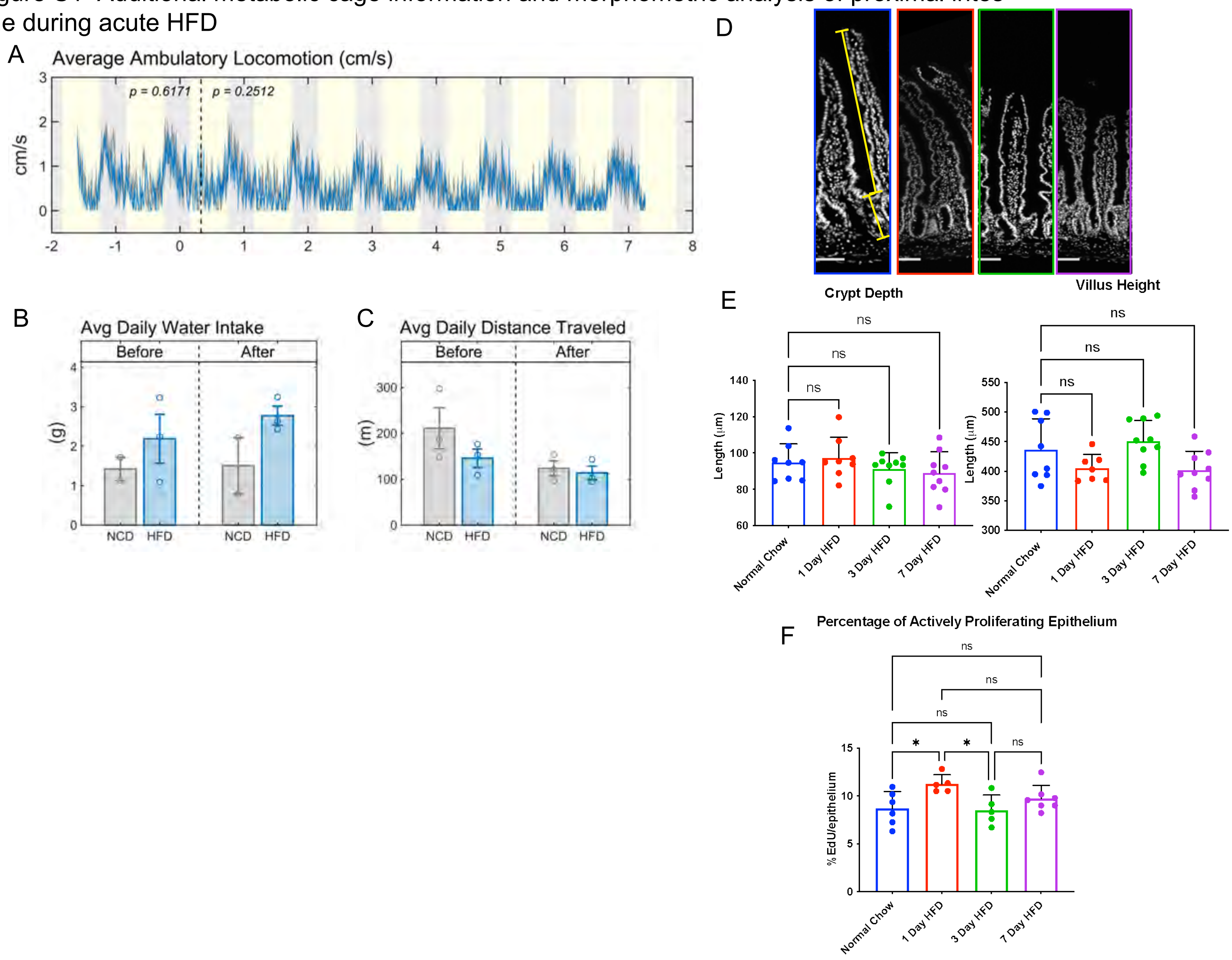

Figure S2- Intestinal Populations of the Proximal Intestine during acute HFD

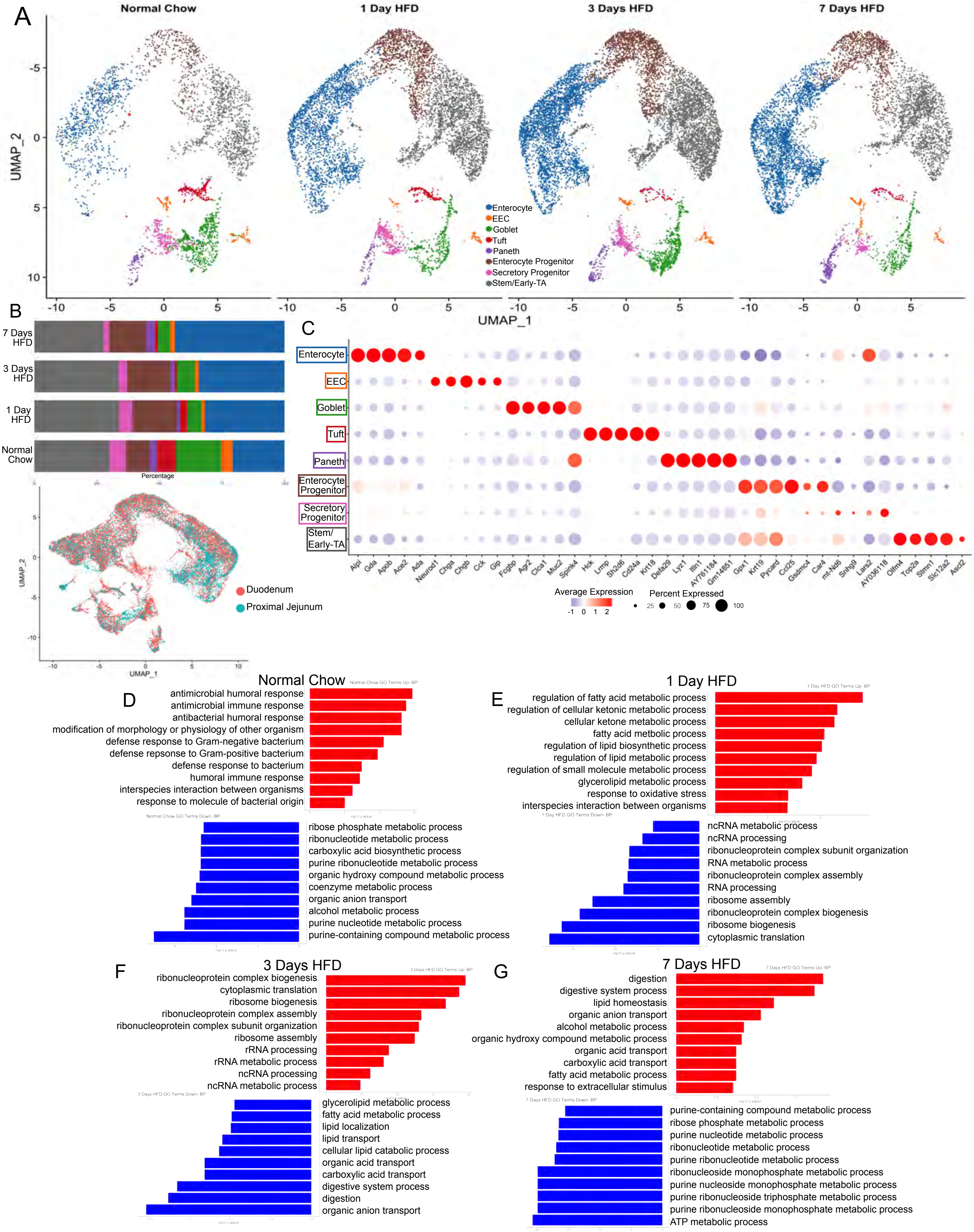

Supplemental Figure 3- Sub-cluster analysis reveals unique responses to acute HFD

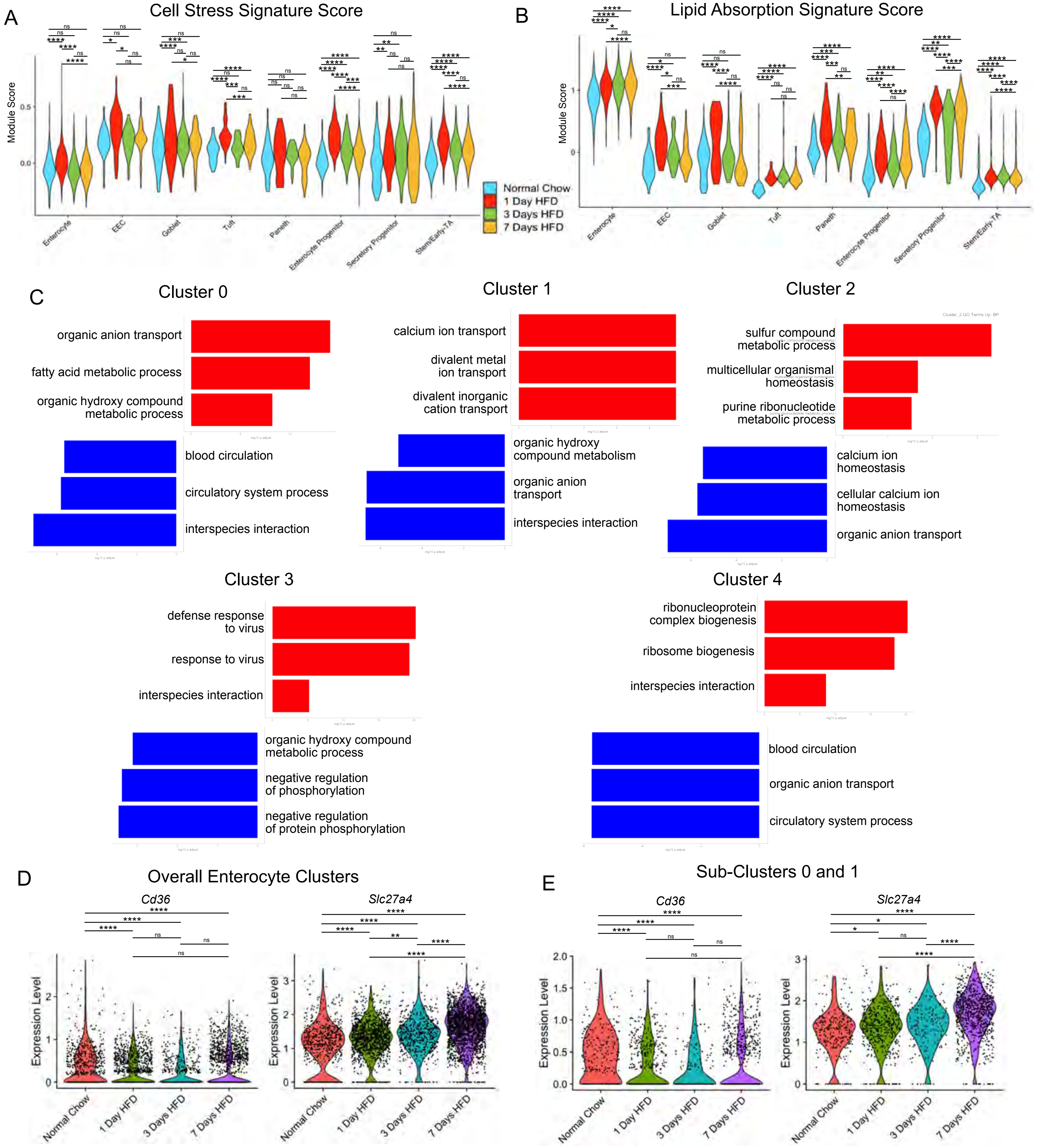

Supplementary Figure 4- *Neurog3* lineage populations enrich for secretory lineages

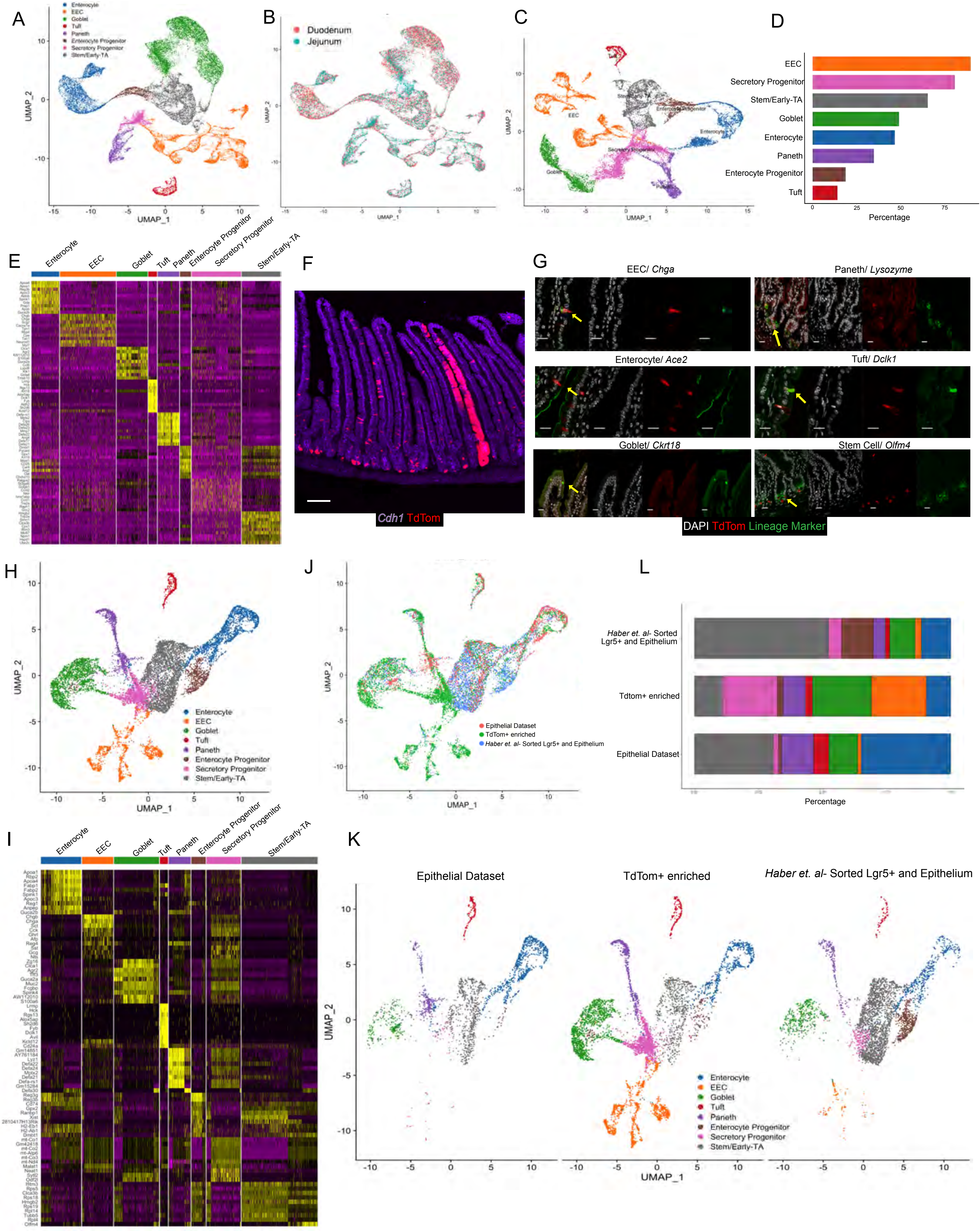

Supplementary Figure 5- EECs Maintain Known Regulatory Role during Acute HFD

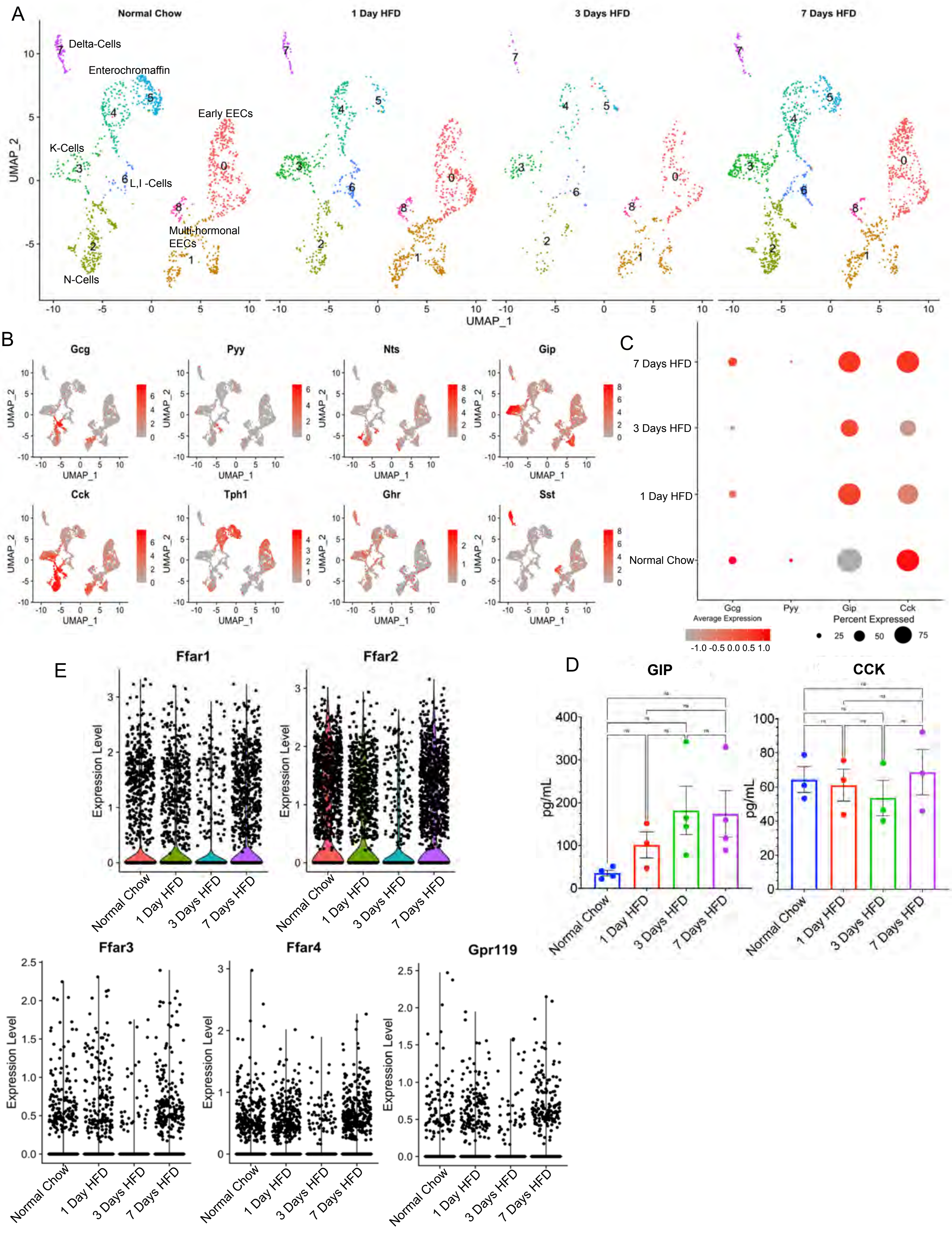

**Supplemental Figure Legends**

Figure S1- Additional Metabolic Cage Information and Morphometric Analysis of Proximal Intestine during Acute HFD

(A) Average ambulatory locomotion of mice fed Normal Chow (gray line) and HFD (blue line) before and after dietary treatment. Yellow color indicates light cycle whereas gray color indicates night cycle. (B) Average daily water intake before and after dietary treatment. NCD = Normal Chow Diet, HFD = High Fat Diet. (C) Average daily total distance traveled in metabolic cages before and after dietary treatment. (D) Representative images of proximal jejunum used for morphometric analysis. Yellow bars indicate representative depth and height measurement. Normal Chow (Blue Border), 1 Day HFD (Red Border), 3 Days HFD (Green Border), 7 Days HFD (Purple Border) with all images showing DAPI (White). Scale bars = 100µm. (E) Quantification of crypt depth (left panel) and villus height (right panel) between dietary conditions; n = 8 for Normal Chow and 1 Day HFD, n = 9 for 3 Days and 7 Days HFD (F) Quantification of 5-ethynyl-2'-deoxyuridine (EdU) incorporation after a 2h pulse between all conditions; n = 6 for Normal Chow, n = 5 for 1 Day and 3 Days HFD, n = 7 for 7 Days HFD (E-F) Error bars are SD. One-Way ANOVA: ns- no significance; p-value * < 0.05

Figure S2- Intestinal populations of the proximal intestine during acute HFD

(A) Integrated analysis of proximal intestine (reference dataset- Normal Chow) sorted for live, epithelial cells and visualized via UMAP split into dietary condition and colorized by cluster identity. (B) Top Panel- Percent proportions of total cells for each cluster over time. Bottom Panel- UMAP distinguishing between cells isolated from duodenum (red) and proximal jejunum (blue) (C) Dot plot depicting average gene expression of top 5 genes expressed for each cluster in proximal jejunum. (D-G) Top 10 Gene Ontology (GO) Terms- Biological Process for overall clusters comparing between each condition in the proximal jejunum. Red bars indicate significantly upregulated whereas blue bars indicate significantly down regulated. X-axis indicates log10 adjusted-p value (cutoff < 0.05).

Figure S3- Sub-cluster Analysis Reveals Unique Responses to Acute HFD

(A) Cell Stress module gene scores for each cluster compared between conditions of the intestinal epithelial dataset. (B) Lipid absorption scores for each cluster of the intestinal epithelial dataset compared between conditions. (C) Top 3 Gene Ontology (GO) Terms- Biological Process for each cluster for the Enterocyte subset comparing between each condition. Red bars indicate significantly upregulated whereas blue bars indicate significantly downregulated. X-axis indicates log10 adjusted-p value (cutoff < 0.05). Extracted from Epithelial dataset. (D) Violin Plots extracted from Enterocyte subset showing gene expression of fatty acid receptors *Cd36* (Cluster of differentiation-36) and *Slc27a4* (Solute carrier Family 27-Member4) split between conditions Normal Chow, 1 Day HFD, 3 Days HFD, 7 Days HFD. (E) Violin Plots depicting gene expression of each cell (black dot) from Enterocyte subset showing gene expression of fatty acid receptors *Cd36* and *Slc27a4* split between conditions Normal Chow, 1 Day HFD, 3 Days HFD, 7 Days HFD. (A-B, C-D) One-Way ANOVA: ns- no significance; p-value * < 0.05, ** < 0.007, *** < 0.0006, **** < 0.0001

Figure S4- *Neurog3* lineage populations enrich for secretory lineages
(A) UMAP of sorted *Neurog3Cre-TdTom* dataset of proximal intestine colored by various cell types. (B) UMAP distinguishing between duodenum and proximal jejunum of TdTom sorted dataset. (C) UMAP of proximal jejunum subset visualizing 8 clusters: EEC (4915 cells), Enterocyte (2437 cells), Enterocyte Progenitor (983 cells), Goblet (2723 cells), Paneth (1890 cells), Secretory Progenitor (4971 cells), Stem/Early-TA Zone (2741 cells), Tuft (766 cells). (D) Percent proportion of each cluster ordered by abundance from top to bottom. (E) Heatmap related to Panel C showing top 10 genes expressed by each cluster. Gradient coloration low (purple) to high (yellow). (F) Immunofluorescence images of proximal intestine from *Neurog3Cre*-tdTom animals. *Cdh1* labels epithelium in purple. Scale bars = 100 μm. (G) Panels of various intestinal epithelial cell types stained in green- Enteroendocrine cell (EEC) stained by Chromogranin-A (*Chga*), Paneth stained by Lysozyme (*Lys*), Enterocyte stained by Angiotensin-converting enzyme 2 (*Ace-2*), Tuft stained by Doublecortin Like Kinase 1 (*Dclk1*), Goblet stained by Cytokeratin-18 (*Ckrt18*), Stem Cells stained by Olfactomedin 4 (*Olfm4*). DAPI counterstains nuclei in white. Scale bars = 20 μm. (H) UMAP colored by cell type after integrated analysis between non-labeled intestinal epithelial cells, sorted TdTomato cells, and *Haber et. al, 2017* dataset. (I) Heatmap related to Panel G of top 10 highly expressed genes for each cell type. (J) UMAP colored by various datasets. (K) UMAP split between each dataset and colored by each cell type. (L) Barplot showing percent abundance of each cell type split between datasets.

Figure S5- EECs maintain known Regulatory Role during Acute HFD
(A) Integrated reference analysis for sorted TdTom+ population (secretory enriched) focusing on enteroendocrine cells (EECs) colored by each cluster and then split into dietary conditions. Total of 8 clusters: 0 (Early EECs, 1234 Cells), 1 (Multi-hormonal EECs, 924 cells), 2 (N-Cells, 722 cells), 3 (K-Cells, 538 cells), 4 (Enterochromaffin-1, 515 cells), 5 (Enterochromaffin-2, 359 cells), 6 (L, I- Cells, 255 cells), 7 (Delta- Cells, 212 cells), 8 (Multi-hormonal EECs, 156 cells); Normal Chow (1448 Cells), 1 Day HFD (1304 cells), 3 Days HFD (502 cells), 7 Days HFD (1661 cells). (B) UMAP feature plots showing gene expression of Glucagon (*Gcg*), Peptide-yy (*Pyy*), Neurotensin (*Nts*), Gastric-Inhibitory Peptide (*Gip*), Cholecystokinin (*Cck*), Tryptophan Hydroxylase 1 (*Tph1*), Ghrelin (*Ghr*), Somatostatin (*Sst*). Colorization is based on normalized expression gradient scale low (gray) to high (red). (C) Average gene expression dot plot for Glucagon (*Gcg*), Peptide-yy (*Pyy*), Gastric-Inhibitory Peptide (*Gip*), and Cholecystokinin (*Cck*) split between dietary conditions. Colorization is based on normalized expression gradient scale low (gray) to high (red). Dot size is based on percentage of cells expressing gene of interest. (D) Enzyme linked immunosorbent assay (ELISA) quantification of GIP and CCK from mouse serum measured in pg/mL. n = 3 or 4 per condition. (E) Violin Plots showing expression of EEC fatty acid receptors *Ffar1, Ffar2, Ffar3, Ffar4* (Free-fatty acid receptor 1-4) and *Gpr119* (G-protein Receptor-119) across conditions Normal Chow, 1 Day HFD, 3 Days HFD, and 7 Days HFD.
